# Fragile streak goals induce pressure responses and an inverted-U performance pattern

**DOI:** 10.64898/2026.02.20.706945

**Authors:** Kagari Yamada, Kazushi Tsutsui, Kazutoshi Kudo

## Abstract

Psychological pressure is thought to relate to performance in an inverted-U pattern, yet evidence is mixed, possibly because manipulations rarely produce high pressure. We induced scalable pressure using a streak goal that resets after a failure in a force-control task. Participants pursued ten consecutive successes (streak goal) or 100 successes irrespective of sequence (total goal). Under the streak goal, heart rate, pupil size, and perceived pressure rose as participants approached their maximum streak; under the total goal, heart rate and pupil size showed little modulation. Performance followed an inverted-U under the streak goal—improving then declining at the maximum streak—whereas the total goal showed no late-stage drop. This dissociation suggests the late-stage decline reflects pressure, not the streak itself. Despite this clear performance decrement, analyses of movement vigor, feedforward/feedback kinematics, and individual differences in pressure responses revealed no consistent systematic signatures at the group level. Fragile streak goals thus provide a multimodal pressure manipulation and a platform for testing mechanisms underlying choking in human motor control.

## Introduction

In situations where high financial rewards or social evaluations depend on an individual’s performance, humans and other nonhuman animals sometimes struggle to perform as intended^1^. This phenomenon, known as “choking under pressure,” is defined as “performance decrements due to psychological pressure”^2^. According to the inverted-U hypothesis^3^, performance initially improves as pressure increases but deteriorates once pressure exceeds an optimal level.

However, empirical support for this inverted U-shaped relationship has been inconsistent^4–7^. One possible reason for this inconsistency is the difficulty in assessing performance across a wide range of arousal states, as humans are rarely pushed into high arousal states in typical cognitive psychology experiments^8^. This limitation may contribute to the lack of clarity regarding the underlying mechanisms of the pressure–performance relationship. For instance, if pressure manipulation as an intervention is insufficient, researchers sample only a limited range of pressure levels, potentially capturing only a partial aspect of the pressure–performance relationship.

In our recent study^9^, we introduced an experimental paradigm in which arousal was progressively induced by setting consecutive success as a goal. When participants aimed to achieve ten consecutive successes, their heart rate, which is a widely used objective measure of arousal often associated with psychological pressure, increased exponentially as the number of successes accumulated. The magnitude of this heart rate increase was substantially greater than that reported in previous studies using more traditional pressure manipulations. However, because heart rate reflects physiological arousal rather than psychological states per se, that study alone could not fully disentangle whether the observed increase in arousal specifically reflected heightened psychological pressure. Inferring a specific psychological state from physiological data alone requires caution, given the issue of reverse inference^10,11^, as other states besides pressure can also influence heart rate^12^.

To address this limitation and rigorously validate the paradigm, the present study incorporated two additional measures. First, we measured pupil size, which reflects emotional arousal linked to sympathetic nervous system activity^13^ and closely tracks trial-by-trial changes in brain state^14^, making it a suitable index for fluctuating arousal under pressure^15,16^. Second, and more critically, we directly assessed participants’ perceived pressure through subjective ratings. This multimodal approach, combining objective physiological indices with subjective reports, allows for a more robust validation of the pressure manipulation, a practice recommended and used in prior pressure research^17–19^.

Using this validated paradigm, we aimed to confirm that the relationship between pressure and performance exhibits the classic inverted U-shape. Participants performed a task requiring precise force control under the goal of achieving ten consecutive successes. Our results confirm that the consecutive success paradigm not only elicits a substantial increase in heart rate, as previously shown, but also leads to a significant increase in pupil size and perceived pressure, thus validating its effectiveness in inducing psychological pressure. Furthermore, we demonstrated that performance followed an inverted U-shaped pattern, improving as pressure increased before deteriorating as it approached the goal.

## Results

Participants were instructed to maintain the cursor within the target area for as long as possible by applying force with their index fingers. This required moving the cursor into the target area as quickly as possible and maintaining its position within the target (Fig. 1A). To induce psychological pressure, we employed the consecutive success paradigm^9^. To dissociate the effects of psychological pressure from those of consecutive success per se, participants were randomly assigned to one of two groups: the Consecutive group or the Total group. The Consecutive group (n = 22) was instructed to achieve 10 consecutive successes, whereas the Total group (n = 22) was instructed to achieve a total of 100 successes, regardless of sequence (Fig. 1B, C). In the Consecutive group, the counter displayed at trial onset reset to zero after any failure, whereas in the Total group the displayed count tracked cumulative successes and therefore remained unchanged after failures. We hypothesized that psychological pressure would increase in the Consecutive group as the number of consecutive successes increased, whereas pressure would remain constant in the Total group, even as the number of consecutive successes increased. By manipulating the overall goals of participants in the experiment, we aimed to assess physiological and behavioral changes induced by psychological pressure in the Consecutive group, and to compare these changes in the Consecutive group with those resulting from consecutive success unrelated to pressure in the Total group.

**Figure 1.**
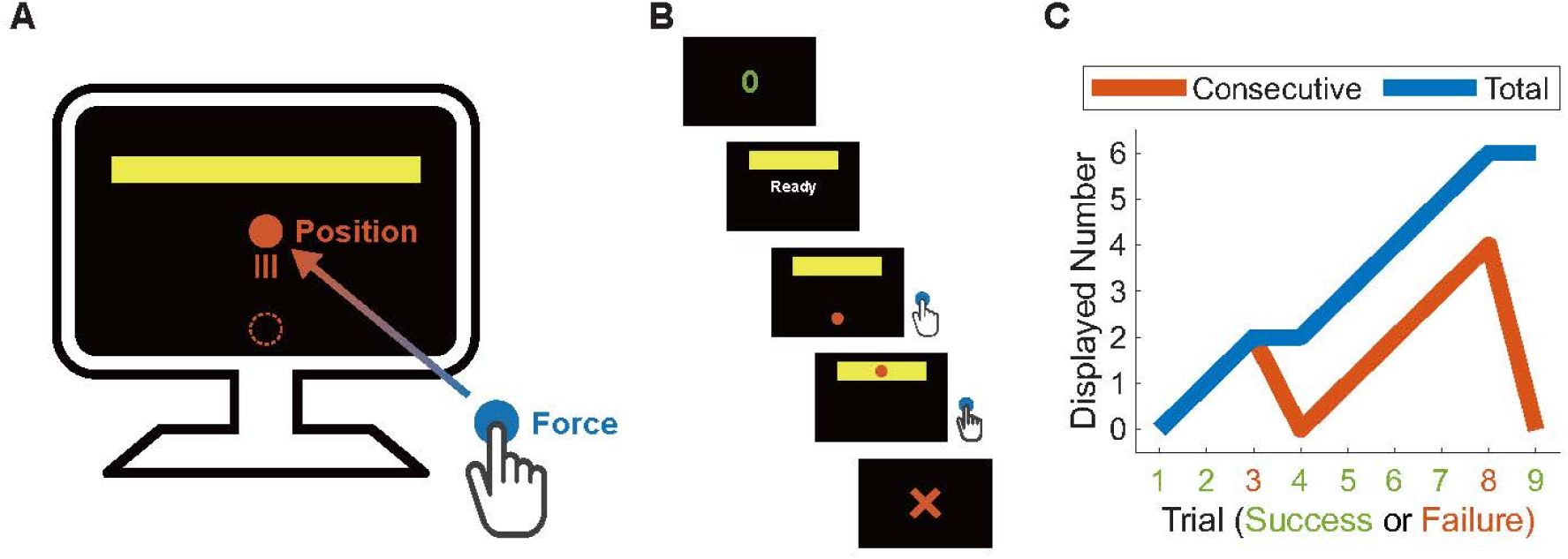
Task and experimental design. (A) Participants performed a task requiring delicate force production. A cursor displayed on the screen moved vertically according to the magnitude of the participant’s exerted force. (B) Sequence of procedures in each trial. First, the participant’s current number of consecutive successes in the Consecutive group or total successes in the Total group was displayed. Then, after a variable interval, participants were required to exert force to move the cursor to the target as quickly as possible and maintain it within the target. Finally, visual feedback of success or failure, determined by the duration the cursor remained within the target, was provided to the participant. (C) Illustration of the number displayed at the beginning of each trial, assuming a hypothetical sequence of successful and unsuccessful trials as shown in the figure. In the Consecutive group, the displayed count reflected the current streak and reset to zero after a failure, whereas in the Total group it reflected cumulative successes and therefore did not reset after failures. Even with the same sequence of outcomes, the numbers displayed at the start of each trial differed between the groups.

### Psychological pressure increased as the number of consecutive successes increased in the Consecutive group only

The occurrence of psychological pressure was assessed using physiological measures of arousal that are sensitive to pressure, namely heart rate^17–20^ and pupil size^15,16^, together with subjective ratings of perceived pressure. To link these measures to progress within a streak in a way that is comparable across participants, we first computed trial-wise averages for each number of consecutive successes and then grouped trials into three within-subject streak-proximity conditions defined relative to each participant’s maximum number of consecutive successes (see Methods for details). In brief, trials were classified as Far, Near, or Max depending on whether they occurred far from, just before, or at the participant’s maximum streak.

Physiological indices confirmed that pressure rose selectively when participants pursued a consecutive-success goal. Heart rate showed a significant main effect of Streak proximity (*F* (1.31, 52.34) = 16.46, *p* < 0.001, *η_G_*^2^ = 0.047) and a Group × Streak proximity interaction (*F* (1.31, 52.34) = 17.44, *p* < 0.001, *η_G_*^2^ = 0.049) (Fig. 2A). In the Consecutive group, heart rate increased monotonically from the Far to Near and Max conditions (*F* (1.23, 24.63) = 19.31, *p* < 0.001, *η_G_*^2^ = 0.137), with all pairwise comparisons significant (Far vs Near: *F* (1, 20) = 16.51, *p* < 0.001, *η_G_*^2^ = 0.095; Near vs Max: *F* (1, 21) = 16.89, *p* < 0.001, *η_G_*^2^ = 0.033). In contrast, heart rate did not vary reliably with streak proximity in the Total group (*F* (1.58, 31.50) = 0.28, *p* = 0.70, *η_G_*^2^ = 0.001).

**Figure 2.**
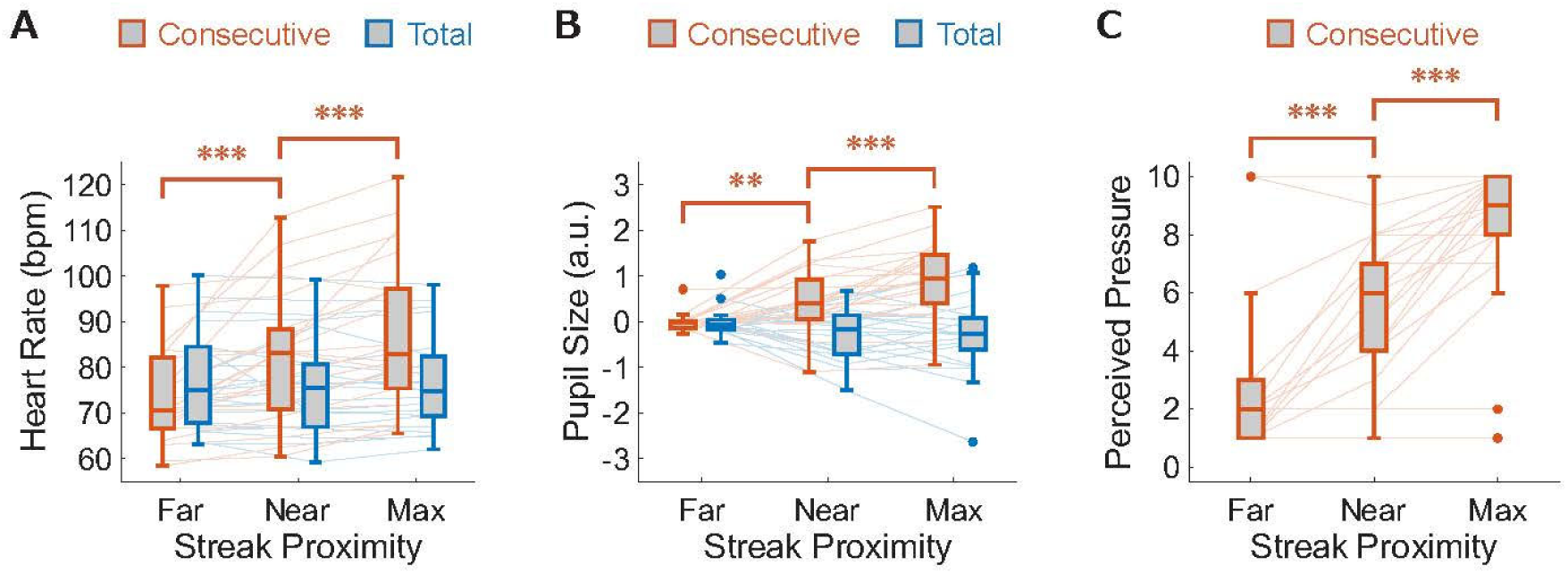
Streak proximity selectively increased pressure indices in the Consecutive group. (A) Heart rate across the three streak-proximity conditions (Far, Near, Max) in the Consecutive and Total groups. (B) Pupil size across the three streak-proximity conditions (Far, Near, Max) in the two groups. (C) Retrospective ratings of perceived pressure across the three streak-proximity conditions (Far, Near, Max) in the Consecutive group. Streak proximity (Far, Near, Max) was defined relative to each participant’s maximum number of consecutive successes (see Methods). In panels A and B, red denotes the Consecutive group and blue denotes the Total group (panel C includes the Consecutive group only). Box-and-whisker plots show the median (center line) and interquartile range (box); whiskers extend to 1.5× the interquartile range and points denote outliers. Thin lines connect individual participants across streak-proximity conditions. Asterisks indicate significant planned pairwise comparisons (\*\**p* < 0.01, \*\*\**p* < 0.001).

A similar pattern was observed for pupil size. Overall pupil size exhibited significant main effects of Group (*F* (1, 38) = 18.23, *p* < 0.001, *η_G_*^2^ = 0.225) and Streak proximity (*F* (1.46, 55.38) = 5.50, *p* < 0.05, *η_G_*^2^ = 0.054), as well as a Group × Streak proximity interaction (*F* (1.46, 55.38) = 15.09, *p* < 0.001, *η_G_*^2^ = 0.135) (Fig. 2B). In the Consecutive group, pupil size increased substantially across streak proximity levels (*F* (1.47, 27.93) = 19.32, *p* < 0.001, *η_G_*^2^ = 0.278), with all pairwise comparisons significant (Far vs Near: *F* (1, 19) = 11.14, *p* < 0.01, *η_G_*^2^ = 0.203; Near vs Max: *F* (1, 20) = 18.86, *p* < 0.001, *η_G_*^2^ = 0.103). In the Total group, pupil size showed no reliable modulation by streak proximity (*F* (1.41, 26.75) = 1.40, *p* = 0.26, *η_G_*^2^ = 0.029).

Since psychological states other than perceived pressure can also influence physiological measures, it is not possible to determine participants’ psychological state solely based on changes in heart rate and pupil size^12^. Previous studies have employed methods that validate pressure manipulation using both subjective and objective measures^17–20^. Therefore, we assessed participants’ perceived pressure using a questionnaire administered after the main session in the Consecutive group and confirmed through subjective measures that their perceived pressure increased with streak proximity (*F* (1.44, 30.31) = 51.7, *p* < 0.001, *η_G_^2^* = 0.440), with all pairwise comparisons significant (Far vs Near: *F* (1, 21) = 37.66, *p* < 0.001, *η_G_^2^* = 0.252; Near vs Max: *F* (1, 21) = 36.89, *p* < 0.001, *η_G_^2^* = 0.213) (Fig. 2C). The convergence of self-reported and physiological measures led us to conclude that the experimental manipulation in this study successfully induced psychological pressure.

### Performance followed an inverted U-shaped pattern as participants approached their maximum streak

The duration for which the cursor remained within the target area was recorded as a score (Fig. 3A). Based on this score, each trial was classified as either a success or a failure. Participants were informed of this scoring criterion and were instructed to maximize their scores. Accordingly, the score can be regarded as a measure of performance in the present experiment.

**Figure 3.**
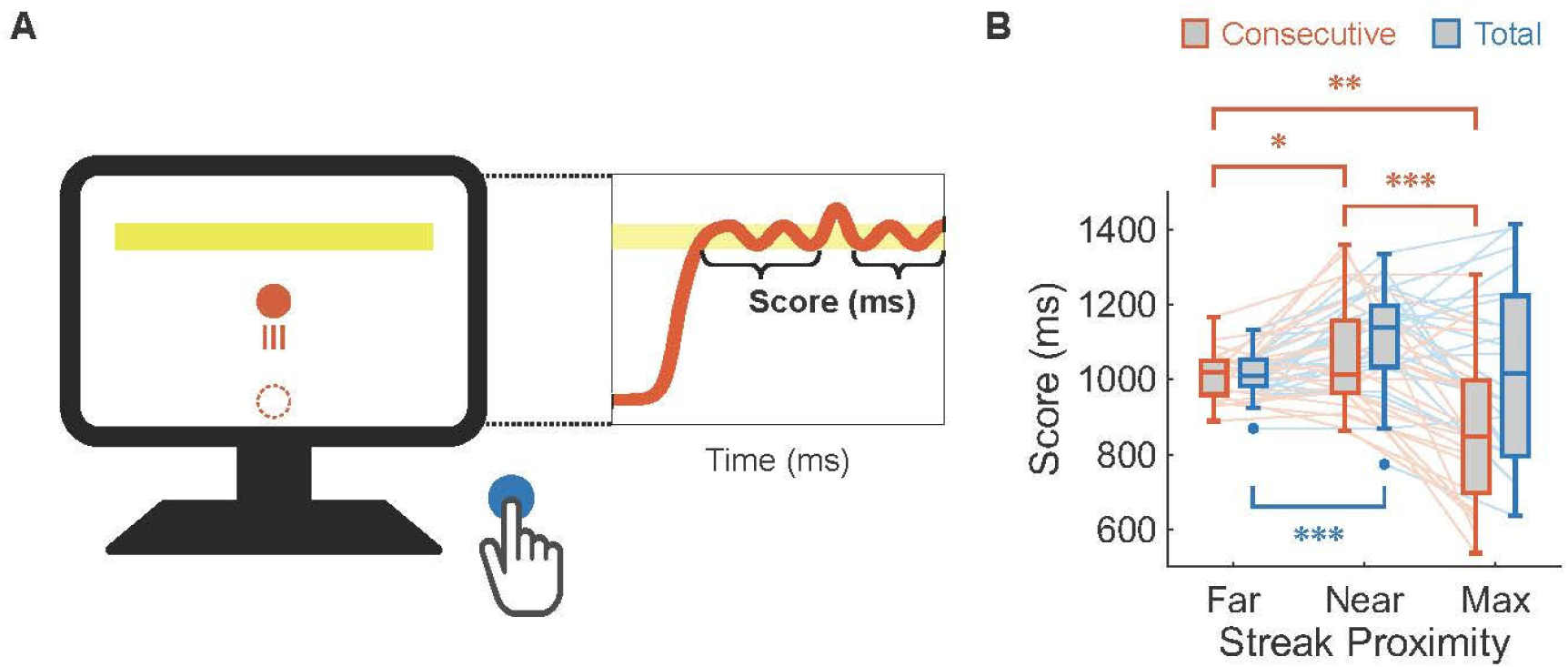
Streak proximity and performance. (A) Schematic of the performance measure. The score was defined as the time (ms) that the cursor remained fully within the target during each trial; trials with score ≥ 1000ms were counted as successes. (B) Performance scores across the three streak-proximity conditions (Far, Near, Max) in the Consecutive (red) and Total (blue) groups. Streak proximity was defined relative to each participant’s maximum number of consecutive successes (see Methods). Red denotes the Consecutive group and blue denotes the Total group. Box-and-whisker plots show the median (center line) and interquartile range (box); whiskers extend to 1.5× the interquartile range and points denote outliers. Thin lines connect individual participants across streak-proximity conditions. Asterisks indicate significant planned pairwise comparisons (\**p* < 0.05, \*\**p* < 0.01, \*\*\**p* < 0.001).

Some studies have reported inverted U-shaped pressure–performance relationships under certain conditions, whereas they have also reported linear or no relationships under other conditions^4–7^. This discrepancy may be attributed to participants’ arousal levels not increasing sufficiently due to a lack of pressure. Because our paradigm produced strong increases in pressure indices as streak proximity increased in the Consecutive group, we examined how performance changed across the three streak-proximity conditions.

A 2 (Group: Consecutive vs. Total) × 3 (Streak proximity: Far, Near, Max) mixed ANOVA on the score revealed significant main effects of Group (*F* (1, 40) = 4.82, *p* < 0.05, *η_G_*^2^ = 0.062) and Streak proximity (*F* (1.51, 60.46) = 13.25, *p* < 0.001, *η_G_*^2^ = 0.129), as well as a Group × Streak proximity interaction (*F* (1.51, 60.46) = 4.62, *p* < 0.05, *η_G_*^2^ = 0.049). Performance varied with streak proximity levels in both groups but with distinct patterns (Fig. 3B).

In the Consecutive group, performance changed markedly across streak proximity levels (*F* (1.64, 32.89) = 12.31, *p* < 0.001, *η_G_*^2^ = 0.222). Scores increased from Far to Near streak proximity (*F* (1, 20) = 4.45, *p* < 0.05, *η_G_*^2^ = 0.072) and then declined at Max streak proximity (*F* (1, 21) = 18.77, *p* < 0.001, *η_G_*^2^ = 0.223). Notably, scores at Max streak proximity were significantly different from those at Far (*F* (1, 20) = 10.12, *p* < 0.01, *η_G_*^2^ = 0.154). Given that pressure indices increased with streak proximity in this group (Fig. 2), this pattern is consistent with an inverted U-shaped relationship between pressure and performance. In the Total group, performance also differed across streak proximity levels (*F* (1.36, 27.22) = 5.18, *p* < 0.05, *η_G_*^2^ = 0.101), driven by improved scores from Far to Near streak proximity (*F* (1, 20) = 28.06, *p* < 0.001, *η_G_*^2^ = 0.276). However, unlike in the Consecutive group, there was no reliable decline in performance from Near to Max streak proximity (*F* (1, 21) = 3.86, *p* = 0.09, *η_G_*^2^ = 0.050), and performance at Max did not differ from Far (*F* (1, 20) = 0.68, *p* = 0.42, *η_G_*^2^ = 0.011).

Thus, increasing streak proximity was associated with performance improvements in both groups, but only when a consecutive-success streak directly determined task termination (Consecutive group) did performance deteriorate at Max streak proximity. Together, these results indicate that the emergence of an inverted U-shaped pressure–performance function is specific to the case in which increasing streak proximity is coupled to a psychologically demanding consecutive-success goal. The absence of a performance drop at Max streak proximity in the Total group suggests that the late-stage decline in the Consecutive group reflects the impact of pressure associated with the streak-based goal structure, rather than the mere fact that a streak happened to lengthen.

### No evidence for increased movement vigor under heightened pressure

It has been well established that human motor behavior changes when larger rewards are at stake—that is, when motivation is heightened—and these changes have been attributed to increases in movement vigor^21^. Increased vigor is associated with reduced reaction times^22,23^, faster movement speeds^24–26^, and greater force production^27,28^. In the present study, we investigated whether the observed behavioral changes could similarly be explained by a change in movement vigor as streak proximity increased.

To this end, we examined whether streak proximity was associated with changes in reaction time, peak movement speed, or force magnitude. For each dependent measure, we averaged trials within the Far, Near, and Max streak-proximity conditions for each participant and submitted these values to a 2 (Group: Consecutive, Total) × 3 (Streak proximity: Far, Near, Max) mixed ANOVA.

There was no evidence that reaction time shortened with streak proximity. For reaction time (Fig. 4A), the mixed ANOVA revealed no significant main effects of Group (*F* (1, 39) = 1.59, *p* = 0.22, *η_G_*^2^ = 0.034) or Streak proximity (*F* (1.29, 50.43) = 0.29, *p* = 0.65, *η_G_*^2^ = 0.001), and no Group × Streak proximity interaction (*F* (1.29, 50.43) = 0.40, *p* = 0.58, *η_G_*^2^ = 0.001). Consistent with this, one-way repeated-measures ANOVAs within each group showed no reliable variation in reaction time across streak-proximity levels in either the Consecutive group (*F* (1.21, 22.95) = 0.54, *p* = 0.50, *η_G_*^2^ = 0.004) or the Total group (*F* (1.36, 27.25) = 0.02, *p* = 0.95, *η_G_*^2^ = 0.000).

**Figure 4.**
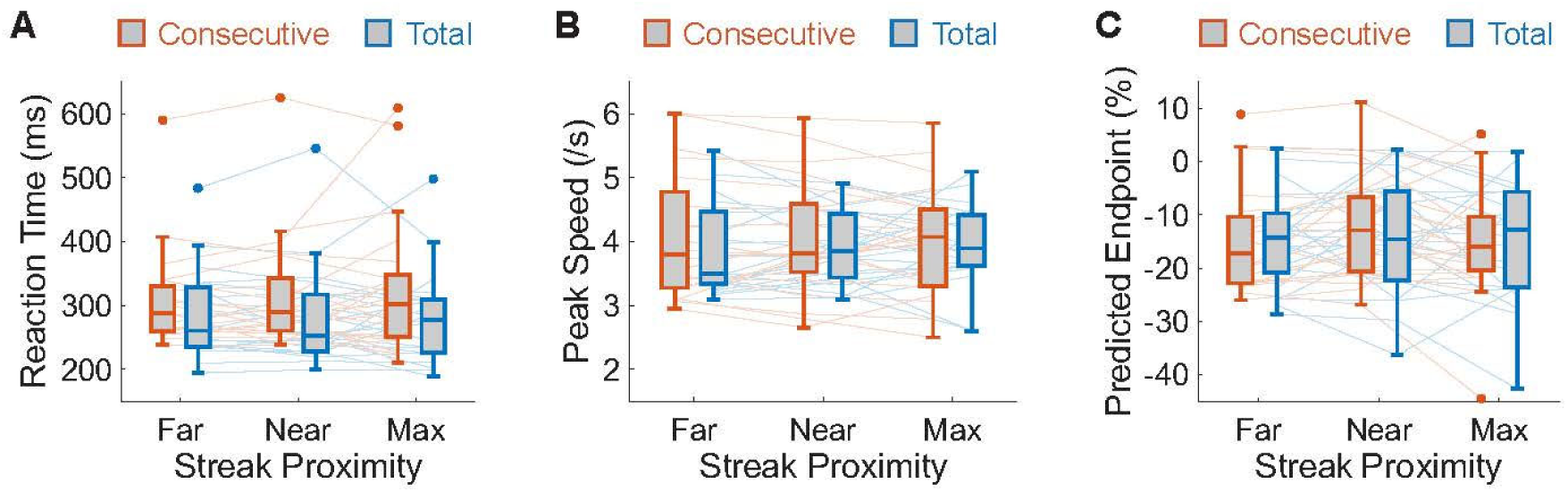
No evidence for increased movement vigor as a function of streak proximity. (A) Reaction time across streak-proximity conditions in the Consecutive (red) and Total (blue) groups. (B) Peak speed across streak-proximity conditions in the two groups. (C) Force-related output index (predicted endpoint relative to the target) across streak-proximity conditions in the two groups. Streak proximity was defined relative to each participant’s maximum number of consecutive successes (see Methods). In panels A–C, red denotes the Consecutive group and blue denotes the Total group. Box-and-whisker plots show the median (center line) and interquartile range (box); whiskers extend to 1.5× the interquartile range and points denote outliers. Thin lines connect individual participants across streak-proximity conditions.

Similarly, streak proximity was not associated with increased movement speed. For peak movement speed (Fig. 4B), the mixed ANOVA again yielded no significant main effects of Group (*F* (1, 39) = 0.41, *p* = 0.53, *η_G_*^2^ = 0.009) or Streak proximity (*F* (1.77, 69.10) = 0.02, *p* = 0.97, *η_G_*^2^ = 0.000), and no Group × Streak proximity interaction (*F* (1.77, 69.10) = 0.54, *p* = 0.56, *η_G_^2^* = 0.001). Within-group analyses confirmed the absence of reliable effects of streak proximity on peak speed in both the Consecutive group (*F* (1.39, 26.34) = 0.27, *p* = 0.69, *η_G_^2^* = 0.001) and the Total group (*F* (1.92, 38.50) = 0.30, *p* = 0.73, *η_G_^2^* = 0.003).

Finally, we examined force-related output, indexed by the predicted endpoint derived from acceleration-phase kinematics (see Methods), relative to the target (Fig. 4C). The mixed ANOVA showed no significant main effects of Group (*F* (1, 39) = 0.04, *p* = 0.84, *η_G_^2^* = 0.001) or Streak proximity (*F* (1.83, 71.55) = 1.25, *p* = 0.29, *η_G_^2^* = 0.007) and no interaction (*F* (1.83, 71.55) = 0.10, *p* = 0.89, *η_G_^2^* = 0.001). One-way ANOVAs within each group likewise showed no reliable modulation of predicted endpoint by streak proximity in either the Consecutive group (*F* (1.20, 22.80) = 1.03, *p* = 0.33, *η_G_^2^* = 0.012) or the Total group (*F* (1.87, 37.36) = 0.33, *p* = 0.71, *η_G_^2^* = 0.004).

Together, these results indicate that the motor changes typically associated with increased movement vigor—shorter reaction times, faster movements, and greater force output—were not elicited as participants approached their maximum streak. Instead, the performance changes associated with increasing streak proximity (and increasing pressure in the Consecutive group) appear to arise from other facets of motor control.

### Exploratory analyses of feedforward and feedback control did not reveal a clear mechanism underlying the inverted U-shaped performance change

Given the null effects on movement-vigor indices, we next conducted exploratory kinematic analyses. Human movement is commonly described as relying on two complementary control processes: feedforward control, which is based on predictions generated before movement initiation, and feedback control, which uses sensory information during movement to correct deviations from those predictions^29^. Here, we asked whether indices intended to capture these processes varied with streak proximity (and, in the Consecutive group, with increasing pressure).

Because the time course of kinematic variables (e.g., velocity) often supports segmentation of goal-directed movements into distinct phases^30–32^, we divided each trial into three phases: an acceleration phase at movement onset, a deceleration phase as the cursor approached the target, and a holding phase after target entry. We reasoned that the acceleration phase would primarily reflect pre-movement predictions, whereas the deceleration and holding phases would be more strongly influenced by visually guided corrections.

To assess whether feedforward accuracy varied with streak proximity, we analyzed the acceleration phase. We estimated where the cursor would have ended if subsequent visually guided corrections had not occurred. Specifically, we mirrored the velocity profile from movement onset to peak cursor speed around the time of peak speed, and then integrated this mirrored profile over time to obtain a predicted endpoint. Predicted error was defined as the absolute relative error between this predicted endpoint and the target center. Predicted error showed no reliable modulation by streak proximity (main effect of Group: *F* (1, 39) = 0.02, *p* = 0.89, *η_G_^2^* = 0.000; main effect of Streak proximity: *F* (1.84, 71.60) = 2.92, *p* = 0.07, *η_G_^2^* = 0.020; Group × Streak proximity interaction: *F* (1.84, 71.60) = 0.08, *p* = 0.91, *η_G_^2^* = 0.001) (Fig. 5A).

**Figure 5.**
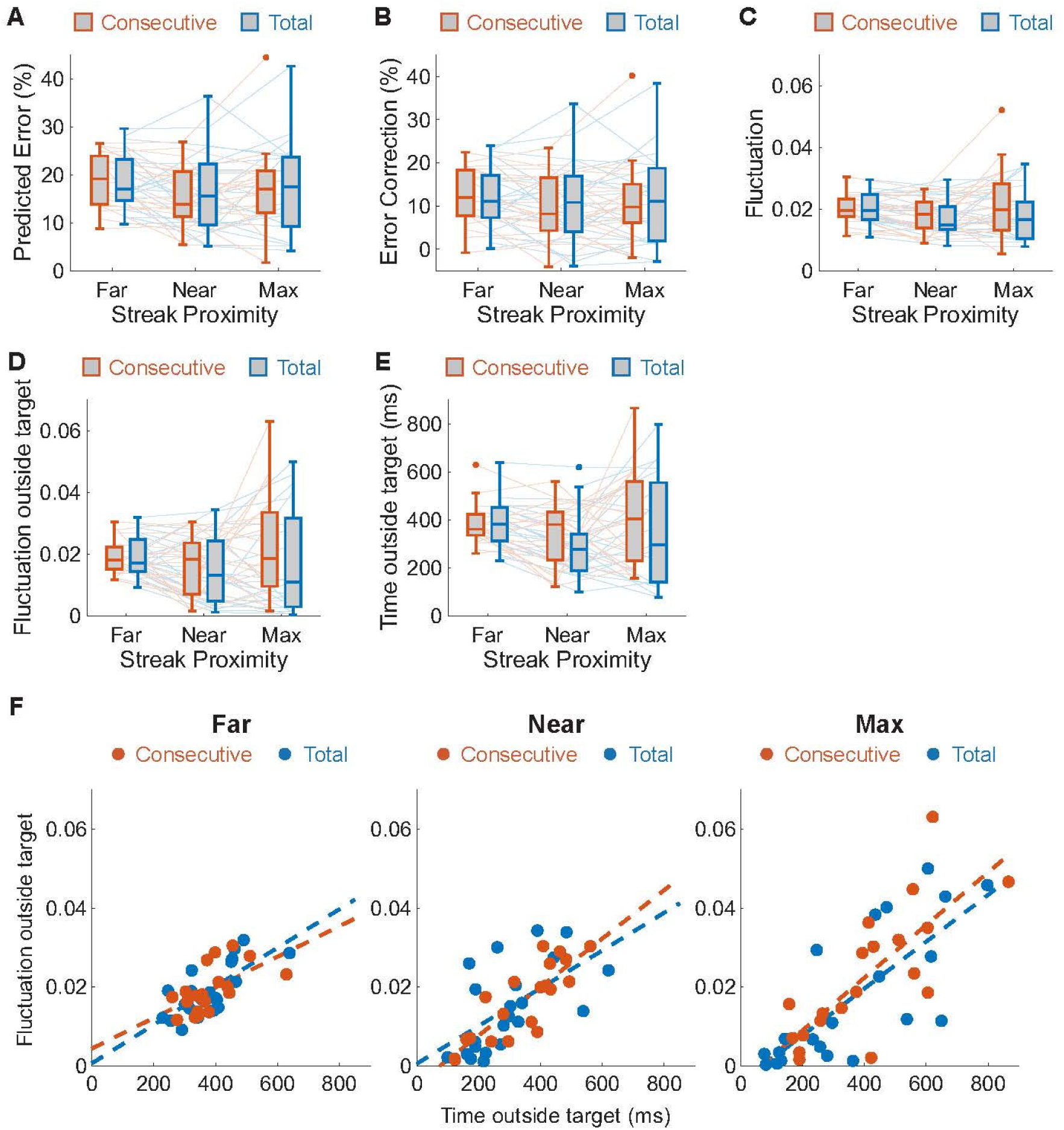
Indices of feedforward and feedback control as a function of streak proximity. (A) Predicted error from the acceleration phase across streak-proximity conditions in the Consecutive (red) and Total (blue) groups. (B) Online correction (error correction) during the deceleration phase across streak-proximity conditions in the two groups. (C) Fluctuation during the holding phase across streak-proximity conditions in the two groups. (D) Fluctuation outside the target (standard deviation of cursor position during out-of-target epochs after initial target entry) across streak-proximity conditions in the two groups. (E) Time outside the target after initial target entry across streak-proximity conditions in the two groups. (F) Relationship between time outside the target and fluctuation outside the target, shown separately for each streak-proximity condition. Streak proximity was defined relative to each participant’s maximum number of consecutive successes (see Methods). In panels A–E, red denotes the Consecutive group and blue denotes the Total group. Box-and-whisker plots show the median (center line) and interquartile range (box); whiskers extend to 1.5× the interquartile range and points denote outliers. Thin lines connect individual participants across streak-proximity conditions. In panel F, each point represents one participant; dashed lines indicate linear fits within each group.

To probe feedback-based correction during the deceleration phase, we computed an online correction (error-correction) index as the difference between predicted error and actual error at the first time point when cursor velocity became negative after peak speed (i.e., predicted error − actual error). Streak proximity did not reliably modulate this correction index (main effect of Group: *F* (1, 39) = 0.00, *p* = 0.98, *η_G_^2^* = 0.000; main effect of Streak proximity: *F* (1.82, 70.90) = 0.91, *p* = 0.40, *η_G_^2^* = 0.006; Group × Streak proximity interaction: *F* (1.82, 70.90) = 0.16, *p* = 0.83, *η_G_^2^* = 0.001) (Fig. 5B). We next examined stability during the holding phase. Fluctuation during the holding phase was defined as the standard deviation of cursor position after target entry. This metric also showed no robust modulation by streak proximity (main effect of Group: *F* (1, 39) = 0.28, *p* = 0.60, *η_G_^2^* = 0.004; main effect of Streak proximity: *F* (1.27, 49.53) = 3.10, *p* = 0.08, *η_G_^2^* = 0.030; Group × Streak proximity interaction: *F* (1.27, 49.53) = 0.55, *p* = 0.50, *η_G_^2^* = 0.006) (Fig. 5C).

Notably, whereas overall task performance (Score) differed across streak-proximity conditions (Fig. 3), the holding-phase fluctuation metric did not. One possible explanation is that the original fluctuation calculation pooled all samples after target entry, potentially mixing epochs in which the cursor was maintained within the target with epochs in which it left the target (hereafter, out-of-target or “error” epochs). If streak proximity influenced behavior primarily during these error epochs, collapsing across both regimes could obscure condition effects.

To test this possibility, we recomputed variability by restricting the standard-deviation calculation to out-of-target epochs only (fluctuation outside target). A 2 (Group) × 3 (Streak proximity) mixed ANOVA provided no evidence for modulation by streak proximity (main effect of Group: *F* (1, 37) = 0.52, *p* = 0.48, *η_G_^2^* = 0.007; main effect of Streak proximity: *F* (1.46, 54.16) = 2.24, *p* = 0.13, *η_G_^2^* = 0.027; Group × Streak proximity interaction: *F* (1.46, 54.16) = 0.47, *p* = 0.57, *η_G_^2^* = 0.006) (Fig. 5D). We next considered whether performance decrements might instead relate to delayed corrective responses to visual feedback. Under a delay account, out-of-target epochs should last longer as participants approach their maximum streak. However, time spent outside the target after initial entry (time outside target) did not show a reliable group-level modulation by streak proximity (main effect of Group: *F* (1, 37) = 0.77, *p* = 0.39, *η_G_^2^* = 0.011; main effect of Streak proximity: *F* (1.27, 46.94) = 3.45, *p* = 0.06, *η_G_^2^* = 0.041; Group × Streak proximity interaction: *F* (1.27, 46.94) = 0.55, *p* = 0.50, *η_G_^2^* = 0.007) (Fig. 5E). Finally, time outside the target and fluctuation outside target were strongly correlated across participants within each streak-proximity condition in both groups (Far: Consecutive: *r* = 0.58, *p* < 0.01, Total: *r* = 0.73, *p* < 0.001; Near: Consecutive: *r* = 0.84, *p* < 0.001, Total: *r* = 0.58, *p* < 0.01; Max: Consecutive: *r* = 0.77, *p* < 0.01, Total: *r* = 0.76, *p* < 0.01) (Fig. 5F). Because this coupling was present at Far, Near, and Max, it is unlikely to provide a mechanism specifically explaining the pressure-related performance drop at Max.

Taken together, these exploratory analyses show that, despite clear increases in pressure indices with streak proximity in the Consecutive group and an inverted U-shaped change in overall performance, the kinematic indices designed to probe feedforward accuracy and feedback-based correction/stability—including alternative indices focusing on out-of-target epochs—did not exhibit robust or systematic modulation by streak proximity in the present ANOVA framework. Thus, while performance followed an inverted U-shaped trajectory as participants approached their maximum streak, the specific movement-level mechanisms underlying this pattern remain unresolved in the current dataset.

### Individual differences in pressure responses and their relation to performance

In the Consecutive group, some participants succeeded in achieving 10 consecutive successful trials, whereas others did not. This variability suggests that while certain individuals demonstrate relative resilience, others experience pronounced performance deterioration under pressure^33^. We examined whether individuals who showed larger within-streak increases in one pressure index also showed larger changes in the other indices, and whether any of these pressure responses predicted how performance changed across streak-proximity levels in the Consecutive group. For each participant, we computed change scores between streak-proximity conditions (Far vs Near, Near vs Max) for perceived pressure, heart rate, pupil size, and performance (score), and then assessed pairwise correlations among these changes.

Despite the clear group-level increases in all three pressure indices (Fig. 2), their covariation across individuals was modest (Fig. 6A, B, C). Changes in perceived pressure were not reliably associated with changes in heart rate (Far vs Near: *r* = 0.02, *p* = 0.94; Near vs Max: *r* = 0.42, *p* = 0.05) or pupil size (Far vs Near: *r* = −0.15, *p* = 0.53; Near vs Max: *r* = 0.36, *p* = 0.11) for either contrast. Likewise, changes in heart rate and pupil size showed only a weak positive tendency to covary (Far vs Near: *r* = 0.15, *p* = 0.54; Near vs Max: *r* = 0.43, *p* = 0.05), which did not reach conventional significance. Thus, participants who reported the largest increases in pressure were not necessarily those who showed the largest physiological changes, and cardiovascular and pupil responses did not align in a simple one-to-one manner.

**Figure 6.**
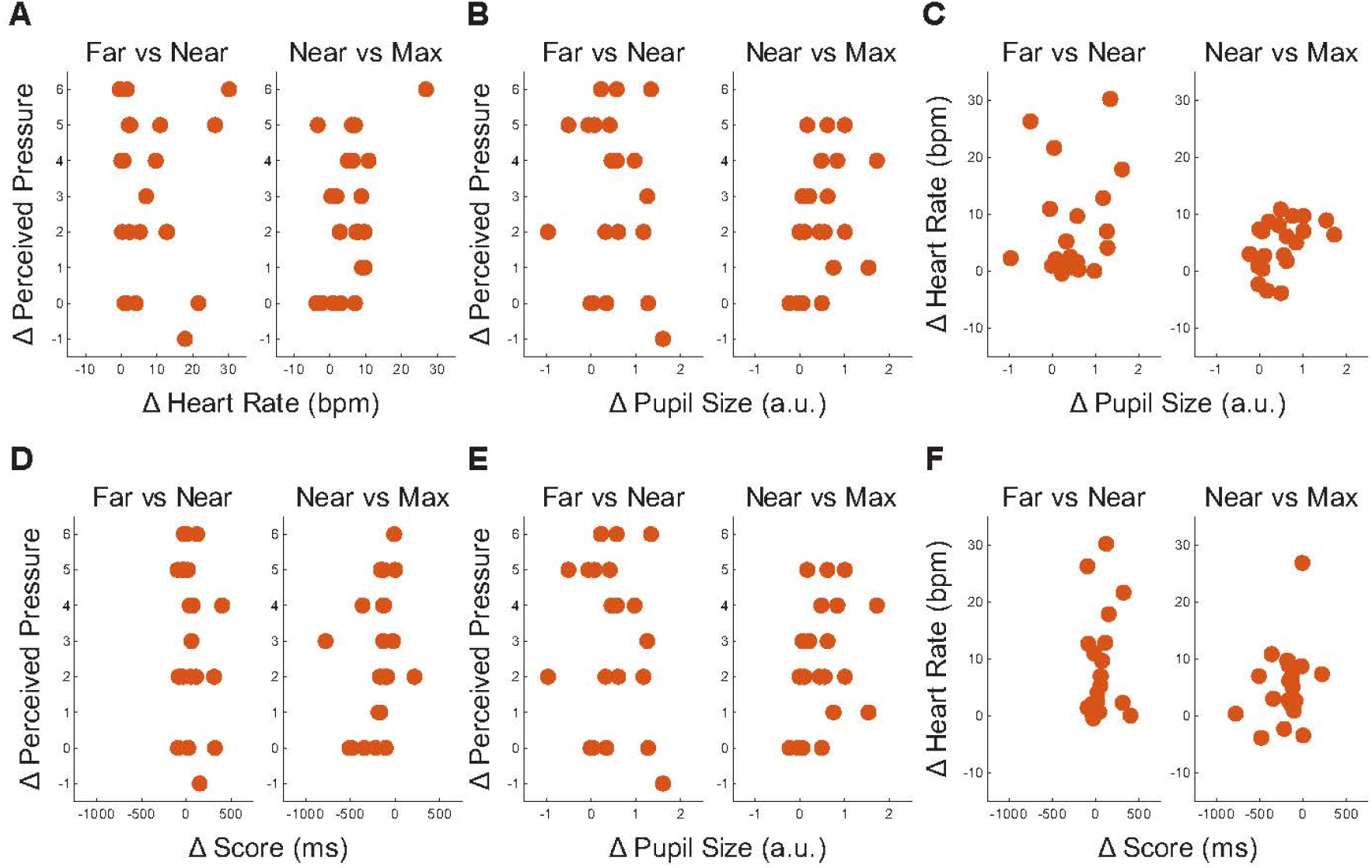
Pairwise associations among pressure indices and between pressure indices and performance. Scatter plots show participant-level change scores (Δ) between streak-proximity conditions in the Consecutive group. (A–C) Associations among pressure indices: (A) Δ perceived pressure vs. Δ heart rate, (B) Δ perceived pressure vs. Δ pupil size, and (C) Δ heart rate vs. Δ pupil size. (D–F) Associations between pressure indices and performance: Δ score plotted against (D) Δ perceived pressure, (E) Δ heart rate, and (F) Δ pupil size. In each panel, the left scatter plot shows changes from Far to Near and the right scatter plot shows changes from Near to Max. Each point represents one participant.

We next asked whether any of these pressure indices predicted how performance changed with increasing streak proximity. Across participants, changes in perceived pressure (Far vs Near: *r* = −0.19, *p* = 0.42; Near vs Max: *r* = 0.31, *p* = 0.17), heart rate (Far vs Near: *r* = 0.10, *p* = 0.67; Near vs Max: *r* = 0.29, *p* = 0.20), and pupil size (Far vs Near: *r* = 0.08, *p* = 0.74; Near vs Max: *r* = 0.13, *p* = 0.56) were all poorly related to changes in score (Fig. 6D, E, F). Individuals with pronounced physiological or subjective responses to pressure did not consistently show larger performance improvements or decrements.

These findings imply that the relationship between pressure and performance is likely influenced by a constellation of factors and cannot be explained solely by a single physiological or subjective measure. This aligns with the view that pressure-induced performance changes are inherently multifaceted^34–37^.

## Discussion

The inverted U hypothesis proposes that performance improves with rising psychological pressure up to an optimal point, but deteriorates once pressure becomes excessive^3^. Empirical support for this nonlinear relationship has been mixed, often appearing only under specific constraints (e.g., task difficulty, incentive framing, instructions, or experimental stage)^4–7^ and potentially because laboratory manipulations seldom push participants into truly high-pressure states^8^. By imposing a “fragile” goal structure—requiring a streak of ten consecutive successes—we elicited strong pressure responses and obtained a clear inverted U-shaped performance curve in a controlled setting, with performance expressed as a function of streak proximity (i.e., proximity to each participant’s max streak).

A central contribution of this work is a more rigorous validation of the consecutive-success paradigm as a manipulation of psychological pressure. In prior work^9^, pressure was inferred primarily from heart rate, leaving room for alternative interpretations because physiological indices alone do not uniquely specify psychological states^10–12^. Here we triangulated pressure using heart rate, pupil size, and perceived pressure ratings—measures that are widely used as indices of pressure, competitive anxiety, and arousal in performance settings^15–20^. All three rose systematically with increasing streak proximity when success had to be maintained as a streak, whereas physiological indices showed little systematic modulation with increasing streak proximity when the goal was to accumulate a fixed number of successes irrespective of sequence. This comparison indicates that it is not repeated success per se, but the coupling between each attempt and the preservation of a streak, that robustly amplifies pressure.

The pressure–performance pattern under the streak-based goal helps clarify why choking is most reliably observed when outcomes feel precarious or structurally fragile. Within the streak-based goal structure, as participants approached the end of a required streak (higher streak proximity), performance improved from low to intermediate streak proximity, consistent with accounts in which moderate pressure can support focus and execution^38–40^. Critically, however, performance declined at high streak proximity—where our physiological and subjective measures indicate that pressure was highest—when success depended on preserving the streak, whereas performance under the total-success goal tended to plateau rather than collapse. This dissociation aligns with the idea that choking is most likely when failures are especially costly or salient—mirroring real-world “one-mistake” contexts such as decisive shots and match points—rather than whenever incentives or repetition are present^4–7,41^. More broadly, this pattern is consistent with the possibility that some null or linear findings in prior laboratory studies reflect manipulations that do not reach sufficiently high-pressure regions of the arousal spectrum^8^.

We next asked whether the performance decline under high pressure could be explained by changes in movement vigor. In many reward and motivation contexts, higher stakes invigorate actions, yielding reduced reaction times^22,23^, faster movement speeds^24–26^, and greater force production^27,28^. Our data provided no evidence for this account: reaction time, peak cursor speed, and force-related indices did not vary reliably with streak proximity. These null effects argue against a simple explanation in which high pressure makes participants “move faster/stronger and therefore fail,” and instead point to other mechanisms—potentially involving changes in attention and anxiety that can alter visuomotor execution without overt invigoration^20,42,43^.

To probe candidate motor-control mechanisms more directly, we conducted exploratory kinematic analyses motivated by distinctions between predictive and sensory-driven control^29,44^. However, the accuracy of feedforward and feedback control did not show consistent modulation by streak proximity at the group level. This stands in contrast to evidence from non-human primates, where high-stakes contexts can yield comparatively systematic behavioral and neural changes^45,46^. One possibility is that, in humans, choking reflects heterogeneous “routes” to failure that average out when analyzed at the group level: some individuals may become distractible, others may over-monitor their movements, and still others may change how they weight sensory evidence versus predictions^2,34–37,47,48^. Consistent with this multifaceted view, correlational analyses of change scores yielded no robust associations among heart rate, pupil size, perceived pressure, and performance, suggesting that pressure-related responses do not necessarily covary across people and may only partially overlap across physiological, subjective, and behavioral channels^39,49,50^. Taken together, these findings suggest that pressure-induced performance changes in humans may not be reducible to a single kinematic mechanism, but can emerge through multiple, partially dissociable routes that vary across individuals—and no consistent pressure-related effects were found in kinematics at the group level. Since the present study does not yet specify the causal computational or neural mechanisms, future work should combine this paradigm with more targeted perturbations that directly test how pressure alters sensory–predictive weighting and control policies—for example, by manipulating visual feedback and error signals^51,52^, and by integrating neural measures that have been implicated in choking under pressure^6,15,46^.

Interestingly, the inverted-U performance pattern observed here contrasts with our previous findings using a similar consecutive-success paradigm^9^. In that study, performance improved monotonically as the streak lengthened, even though the goal structure elicited an exponential increase in heart rate comparable to the profound arousal observed in the present study. This discrepancy—where the exact same pressure-inducing framework yields either a linear improvement or a late-stage collapse—suggests that the relationship between pressure and performance is highly task-dependent. Several theoretical and empirical accounts propose that task difficulty dictates whether this relationship is linear or follows an inverted-U curve^3,53,54^. However, the average success rate in the previous study^9^ (51.1 ± 10.1%) and that of the Consecutive group in the present study (47.4 ± 10.8%) did not differ substantially. Thus, it is likely that factors other than overall task difficulty determine how pressure shapes performance trajectories. Because our exploratory kinematic analyses did not pinpoint a specific mechanism, identifying the critical variables that govern the shape of the pressure–performance relationship remains an important avenue for future research. In sum, requiring a streak of consecutive successes provides a strong and scalable way to induce multimodal pressure in the laboratory, extending earlier demonstrations based on heart rate alone^9^ and generalizing the manipulation to a visuomotor task. This manipulation yields a textbook inverted U-shaped relationship between performance and streak proximity, consistent with an inverted-U pressure–performance account, but neither movement vigor measures nor the present kinematic indices revealed consistent streak-proximity–dependent signatures, and individual differences did not align across physiological, subjective, and behavioral measures. Ultimately, by providing a method to easily induce substantial psychological pressure accompanied by subjective and physiological changes in a controlled environment, this study offers a valuable platform for elucidating the effects of pressure on motor control and its underlying mechanisms.

## Methods

### Participants

44 men (mean age ± SD = 25.7 ± 10.0 years, age range: 18–60 years) participated in this study. According to self-reports, four participants were left-handed. All participants had normal or corrected-to-normal vision. None reported a history of cardiac disease. This study was approved by the Ethical Review Committee for Experimental Research Involving Human Subjects, Graduate School of Arts and Sciences, University of Tokyo. The target sample size (n = 44) was determined a priori based on our previous work using the consecutive-success paradigm^9^, which showed robust, progressive increases in heart rate as consecutive successes accumulated, and to allow balanced assignment to the Consecutive and Total groups (n = 22 each). All participants provided written informed consent before participating in the study.

### Apparatus

All visual stimuli were controlled using a computer (Dell, XPS) and displayed on a monitor (I-O DATA, KH2500V-ZX2,1920×1080 pixels, refresh rate: 240 Hz). The participants sat in front of a desk with a monitor and were instructed to place the elbow of their dominant hand on an armrest, with their wrist on a box, and their index finger on a pressure sensor (TEAC, TU-QR(T)500N-G). Their non-dominant hand was instructed to rest in a relaxed manner, with their legs on the same side.

The force exerted by the participants was converted into a voltage signal by the pressure sensor. The voltage signal was transmitted to a data acquisition device (National Instruments, USB-6218 (BNC)) via a digital indicator (TEAC, TD-700T) and digitally recorded at a sampling frequency of 1000 Hz on a computer running the MATLAB Data Acquisition Toolbox.

Heart rates were measured using an electrocardiographic sensor (Delsys, Delsys Trigno EKG Biofeedback Sensor). Two pre-lubricated disposable ECG electrodes (Nihon Kohden, Dispo Electrode M Vitrode) were attached to each participant. The measured data were digitally recorded at a sampling frequency of 519 Hz using an EMGworks Acquisition system (Delsys). A trigger (Delsys, Trigger Module SP-U02) was used to control the ECG sensor using a computer.

Pupil size was measured with an Eyelink 1000 plus (SR Research) controlled by a computer (Dell, OptiPlex) and recorded at 500 Hz. To improve the quality of the pupil size data, head movement was minimized using a chinrest, and calibration was performed at the start of each block.

### Procedure

Participants were randomly assigned to one of two groups, Consecutive or Total. The Consecutive group (22 participants, mean age ± SD = 25.1 ± 8.23 years, age range: 19–53 years) was told that the experiment would end after 10 consecutive successes, whereas the Total group (22 participants, mean age ± SD = 26.3 ± 11.4 years, age range: 18–60 years) was told that the experiment would end after 100 successes in total.

After the maximum force exerted on their index fingers was measured, the participants performed a training session (2 blocks). In the training session, success resulted in a 1% of the height of the monitor decrease in the target size for the subsequent trial, while failure led to a 2% of the height of the monitor increase in the target size. The first block started with the target size of 20% of the height of the monitor. The second block started with the target size at the end of the first block. The target size in the main session was calculated by averaging all reversals in the second block in the training session.

After the training session, the main session began. A block consisted of 50 trials. If the 50th trial was successful, the trials in that block continued until failure. Participants took a five-minute break between blocks. The experiment continued until participants achieved 10 consecutive successes in the Consecutive group and until they reached a total of 100 successful trials in the Total group. However, if they could not, the experiment was terminated 90 min after the start of the experiment. The total number of trials for each participant varied depending on their performance and which of the two stopping criteria was met first. The resulting trial counts for each participant are detailed in Tables S1 and S2. Participants were informed of these conditions beforehand.

After the main session, we administered a questionnaire to participants in the Consecutive group in which they retrospectively rated perceived pressure. Retrospective assessment was adopted to avoid interrupting the task and altering participants’ attentional focus or evaluative context during the experiment. Participants rated perceived pressure using a VAS ranging from 1 (“felt no pressure”) to 10 (“felt a lot of pressure”). Participants in the Total group were not aware of the number of consecutive successes, so it would be difficult to rate perceived pressure as a function of streak proximity. Therefore, perceived pressure was not investigated in the Total group.

### Task

First, the current number of consecutive successes was presented in the Consecutive group, and the total number of successes at that point was presented in the Total group. After that, the word “Ready” appeared at the center of the monitor, and a target was displayed. The lower edge of the target was positioned 70% of the height of the monitor from the bottom of the monitor and the target size was based on performance during training session. During this phase, participants were instructed not to exert any force on the pressure sensor and the sensors were calibrated. A cursor would be positioned at the center of the target when 10% of their maximum force was exerted. Then “Ready” disappeared, and the cursor appeared 10% of the height of the monitor from the bottom of the monitor. Participants were instructed to move the cursor representing the force exerted by them into the target as quickly as possible upon the cursor appearance, and to hold the cursor within the target until it disappeared. Real-time feedback of the force exerted by participants was provided as the cursor moving on the monitor. The cursor disappeared 2000ms after its appearance. A trial was classified as successful if the cursor remained within the target for at least 1000ms within this interval. Finally, participants received visual feedback on whether they could keep their cursor within the target for more than 1000ms. All visual stimuli and programs were created using MATLAB.

### Analysis Heart rate

All heart rate data were analyzed using a custom-made MATLAB program. The difference between the heart rate in each trial and the average heart rate in the second block in the training session was obtained as the heart rate data for each trial. To remove the effects of the task outcome, the data when the visual feedback of result in each trial was displayed were excluded, that is, the data from the presentation of the number of consecutive successes or the total number of successes just before the presentation of the trial results were analyzed. Data from 5 trials by one participant were excluded because of the failure to acquire heart rate data.

### Pupil Size

All pupil size data were analyzed using a custom-made MATLAB program. First, we discarded 50 ms of pupil size data before and after a blink event. We then interpolated over the discarded data using a piecewise cubic Hermite interpolating polynomial (Matlab *pchip.m* function). After the interpolation, the data were smoothed (*smoothdata.m* function). To remove the effects of gaze position during force exertion and the task outcome, the data during force exertion of participants and the data when the visual feedback of result in each trial was displayed were excluded, that is, the data from the presentation of the number of consecutive successes or the total number of successes just before their force exertion were analyzed. To control for individual differences in baseline levels and dynamic ranges, trial-wise data for pupil size were z-score normalized within each participant before aggregation. Data from one participant were excluded because of the failure to acquire the pupil size data.

### Behavioral Data

All behavioral data were analyzed using a custom-made MATLAB program. The voltage data sampled at 1000 Hz were converted so that the cursor would be positioned at the center of the target when the participant exerted 10% of their maximum force. The total time that the cursor was completely within the target in each trial was calculated as the score. If the score was 1000 ms or more, it was judged a success, otherwise, it was judged a failure.

Following the approach of previous research^45^, we conducted a detailed analysis of cursor movement. We divided the exertion of force in the task into three phases: an acceleration phase at the beginning of the movement, a deceleration phase when the cursor approaches the target, and a holding phase within the target. To calculate predicted errors from the acceleration phase, we first found the time of peak speed of the cursor. We then took the velocity profile from the appearance of the cursor until this time and reflected it around the time of peak speed. The resulting symmetric velocity profile was then integrated to find the displacement from the position of the appearance of the cursor, which meant a predicted endpoint based on data before the time of peak speed. The absolute value of the relative error between the predicted endpoint and the center of the target was calculated as a “predicted error from the acceleration phase”. For correction during the deceleration phase, we wanted not only a predicted endpoint but also an actual endpoint. We identified the time point at which the cursor velocity became negative following its peak, and defined the cursor position at that moment as the actual endpoint. The absolute error between the actual endpoint and the target center was defined as the actual error at the end of the deceleration phase. By subtracting this value from the predicted error from the acceleration phase, we quantified error correction during deceleration. In addition, the standard deviation of the cursor position after the cursor entered the target was calculated as a “fluctuation during the holding phase”. Reaction time for each trial was estimated based on cursor velocity data from the onset of the movement cue to the time of peak velocity. Assuming a single change point in the velocity profile, *ischange.m* function was used to detect the point that minimized the total cost function^55^.

Trials that met any of the following criteria were excluded from the detailed analysis of these cursor movements: “score was 0”, “force was already being exerted when the cursor appeared”, or “the speed of the cursor did not become negative”. As more than half (148 out of 263 trials) of the data from one participant in the Consecutive group met the exclusion criteria, all of that participant’s data was excluded from the detailed analysis. For all of the other participants, the number of trials that met the exclusion criteria was 42 out of 4382 trials (1.0%) in the Consecutive group and 26 out of 4367 trials (0.6%) in the Total group.

### Statistical Analysis

To examine how streak proximity related to physiological, behavioral, and kinematic measures (and, in the Consecutive group, the concomitant increase in psychological pressure), we transformed the trial-wise data into three within-participant streak-proximity conditions. For each participant, we identified their maximum number of consecutive successes across the experiment. Trials were then binned into: (i) Far streak proximity, comprising trials from the beginning of a run up to three successes below the individual maximum; (ii) Near streak proximity, comprising trials at two and one successes below the maximum; and (iii) Max streak proximity, comprising trials at the participant’s maximum consecutive successes. For the Total group, the maximum number of consecutive successes was truncated at nine to match the range of the Consecutive group. Within each bin, we averaged the dependent variables across trials, yielding three observations (Far, Near, Max) per participant and measure. This asymmetric binning (broader “Far” and finer bins near the maximum) was motivated by prior work showing that heart rate increases approximately exponentially as consecutive successes accumulate, such that the steepest physiological changes are expected near the end of a streak^9^.

For physiological and kinematic measures, these averaged values were submitted to mixed-design analyses of variance (ANOVAs) with a between-participant factor Group (Consecutive vs. Total) and a within-participant factor Streak proximity (Far, Near, Max). Sphericity for the within-participant factor was evaluated using the Greenhouse–Geisser epsilon. When the sphericity assumption was violated, we report Greenhouse–Geisser–corrected degrees of freedom and p-values. We report F statistics (with corresponding degrees of freedom), p-values, and generalized eta squared (η_G_^2^).

Perceived pressure ratings in the Consecutive group were analyzed using a one-way repeated-measures ANOVA with the within-participant factor Streak proximity (Far, Near, Max), defined in the same way as for the physiological and kinematic measures. Planned pairwise comparisons between streak-proximity levels were conducted within each group. For consistency with the ANOVA reporting, pairwise tests are reported as F statistics with 1 numerator degree of freedom (equivalent to squaring the paired-sample t statistic), and p-values were adjusted using the Benjamini–Hochberg false discovery rate (FDR) procedure. For all analyses, the significance level was set at α = 0.05.

## Author contributions

Conceptualization and methodology, K.Y. and K.K.; investigation, formal analysis, data curation, visualization, and writing – original draft, K.Y.; writing—review & editing, K.T. and K.K.; funding acquisition, K.Y., K.T. and K.K.; resources, K.K.; supervision, K.K. All authors read and approved the final manuscript.

## Acknowledgments

This work was supported by KAKENHI (22K17673, 24K02825, 24H01434, 25H01237, and 25KJ1145) from the Japan Society for the Promotion of Science (JSPS), and Strategic Basic Research Programs ACT-X (JPMJAX24LA) from the Japan Science and Technology Agency, and Kubota endowed fund from Kubota Co., Ltd. K.Y. was supported by JSPS as a JSPS Research Fellow.

