## Supplementary information for "Fragile streak goals induce pressure responses and an inverted-U performance pattern"

Table 1. Number of trials in each consecutive success for each participant in the Consecutive group.

| Participant | Number of Consecutive Success | | | | | | | | | | |
|---|---|---|---|---|---|---|---|---|---|---|---|
| | 0 | 1 | 2 | 3 | 4 | 5 | 6 | 7 | 8 | 9 | Total |
| 1 | 119 | 40 | 23 | 13 | 7 | 5 | 2 | 0 | 0 | 0 | 209 |
| 2 | 170 | 52 | 22 | 9 | 4 | 2 | 1 | 0 | 0 | 0 | 260 |
| 3 | 10 | 5 | 4 | 2 | 2 | 2 | 1 | 1 | 1 | 1 | 29 |
| 4 | 37 | 18 | 11 | 3 | 2 | 2 | 2 | 1 | 1 | 1 | 78 |
| 5 | 76 | 30 | 17 | 10 | 4 | 3 | 3 | 3 | 2 | 2 | 150 |
| 6 | 27 | 12 | 9 | 6 | 6 | 5 | 3 | 2 | 2 | 2 | 74 |
| 7 | 116 | 50 | 32 | 19 | 10 | 9 | 7 | 4 | 3 | 1 | 251 |
| 8 | 156 | 56 | 26 | 9 | 3 | 0 | 0 | 0 | 0 | 0 | 250 |
| 9 | 164 | 66 | 19 | 3 | 0 | 0 | 0 | 0 | 0 | 0 | 252 |
| 10 | 119 | 64 | 33 | 23 | 12 | 6 | 4 | 0 | 0 | 0 | 261 |
| 11 | 130 | 56 | 34 | 20 | 6 | 4 | 2 | 0 | 0 | 0 | 252 |
| 12 | 172 | 37 | 18 | 12 | 9 | 5 | 4 | 3 | 3 | 0 | 263 |
| 13 | 125 | 51 | 29 | 17 | 12 | 9 | 5 | 4 | 3 | 2 | 257 |
| 14 | 100 | 36 | 21 | 13 | 7 | 5 | 3 | 3 | 3 | 3 | 194 |
| 15 | 95 | 52 | 33 | 19 | 10 | 8 | 1 | 1 | 1 | 1 | 221 |
| 16 | 152 | 51 | 30 | 15 | 4 | 0 | 0 | 0 | 0 | 0 | 252 |
| 17 | 130 | 66 | 27 | 13 | 9 | 4 | 2 | 1 | 1 | 1 | 254 |
| 18 | 52 | 28 | 18 | 10 | 8 | 5 | 3 | 3 | 1 | 1 | 129 |
| 19 | 159 | 51 | 26 | 7 | 3 | 2 | 2 | 1 | 0 | 0 | 251 |
| 20 | 154 | 54 | 27 | 14 | 4 | 1 | 0 | 0 | 0 | 0 | 254 |
| 21 | 113 | 58 | 37 | 21 | 13 | 4 | 4 | 2 | 0 | 0 | 252 |
| 22 | 186 | 47 | 16 | 2 | 1 | 0 | 0 | 0 | 0 | 0 | 252 |
| Ave. | 116.5 | 44.5 | 23.3 | 11.8 | 6.2 | 3.7 | 2.2 | 1.3 | 1.0 | 0.7 | 211.1 |

Note. Each column shows the number of trials in each consecutive success, and each row shows the number of trials for each participant.

Table 2. Number of trials in each consecutive success for each participant in the Total group.

| Participant | Number of Consecutive Success | | | | | | | | | | |
|---|---|---|---|---|---|---|---|---|---|---|---|
| | 0 | 1 | 2 | 3 | 4 | 5 | 6 | 7 | 8 | 9 | Total |
| 1 | 155 | 59 | 24 | 11 | 4 | 0 | 0 | 0 | 0 | 0 | 253 |
| 2 | 31 | 20 | 11 | 8 | 7 | 6 | 6 | 4 | 3 | 3 | 130 |
| 3 | 102 | 38 | 19 | 13 | 8 | 6 | 5 | 4 | 2 | 2 | 201 |
| 4 | 45 | 30 | 21 | 15 | 10 | 8 | 5 | 2 | 2 | 2 | 144 |
| 5 | 117 | 35 | 22 | 14 | 9 | 8 | 4 | 3 | 2 | 2 | 216 |
| 6 | 62 | 35 | 20 | 13 | 8 | 7 | 5 | 4 | 2 | 2 | 161 |
| 7 | 164 | 47 | 23 | 10 | 7 | 3 | 1 | 0 | 0 | 0 | 255 |
| 8 | 88 | 48 | 27 | 11 | 7 | 2 | 2 | 1 | 1 | 0 | 187 |
| 9 | 33 | 26 | 20 | 14 | 10 | 8 | 6 | 4 | 2 | 2 | 132 |
| 10 | 108 | 43 | 21 | 12 | 6 | 5 | 2 | 2 | 2 | 1 | 207 |
| 11 | 66 | 35 | 18 | 12 | 10 | 7 | 5 | 2 | 2 | 2 | 165 |
| 12 | 124 | 49 | 21 | 13 | 9 | 4 | 1 | 1 | 1 | 0 | 223 |
| 13 | 104 | 40 | 19 | 13 | 6 | 5 | 4 | 4 | 4 | 2 | 203 |
| 14 | 119 | 52 | 23 | 13 | 6 | 3 | 2 | 0 | 0 | 0 | 218 |
| 15 | 198 | 39 | 10 | 3 | 0 | 0 | 0 | 0 | 0 | 0 | 250 |
| 16 | 120 | 51 | 27 | 11 | 6 | 2 | 2 | 0 | 0 | 0 | 219 |
| 17 | 84 | 44 | 23 | 13 | 6 | 4 | 3 | 2 | 2 | 1 | 183 |
| 18 | 62 | 40 | 26 | 14 | 9 | 4 | 3 | 2 | 1 | 0 | 161 |
| 19 | 104 | 42 | 25 | 10 | 5 | 4 | 3 | 3 | 3 | 2 | 203 |
| 20 | 70 | 35 | 23 | 14 | 9 | 6 | 3 | 2 | 2 | 2 | 169 |
| 21 | 126 | 56 | 21 | 13 | 5 | 2 | 1 | 1 | 0 | 0 | 225 |
| 22 | 126 | 49 | 18 | 10 | 4 | 2 | 1 | 1 | 1 | 0 | 212 |
| Ave. | 100.4 | 41.5 | 21.0 | 11.8 | 6.9 | 4.4 | 2.9 | 1.9 | 1.5 | 1.0 | 196.2 |

Note. Each column shows the number of trials in each consecutive success, and each row shows the number of trials for each participant.
